# Aggressive responses to rivals depend on the interaction between the vocal traits of territory-holder and mimicked intruders in a miniature tropical frog

**DOI:** 10.1101/2024.04.16.589714

**Authors:** Matías I. Muñoz, Nicolás Camargo-Rodriguez, Wouter Halfwerk

## Abstract

Territory defence is an important aspect of animal behaviour, and tightly linked to rival assessment and species recognition processes. An important component of these two processes is signalling, and animals must decide how to respond to a challenge based on the signals they perceive from their rivals. Here we manipulated the note repetition rate of rocket frog calls to simulate rivals with high and low vocal performances, and tested the aggressive responses of males in the field. We found that the probability of aggression depended on the interaction between the stimulus treatment and the peak frequency of the territorial males. Low-frequency calling males were more likely to respond aggressively to the fast call, while high-frequency calling males showed a higher probability of aggression towards the slow call. Frogs that responded aggressively approached the slow call 2.4-times faster than the fast call. Low-frequency calling males also increased their peak frequencies in response to the playbacks, while high-frequency calling males did not modify their vocalizations. Body weight was not correlated with the peak frequency or the inter-note interval of males. Peak frequency and amplitude (loudness) are positively interrelated in frogs, and are potential indicators of the motivational state of the signaller. We argue that low-frequency males were not singing at their maximum capacity, and they increased their vocal output when challenged by a rival, especially when the rival was a high-performance individual. We show that rocket frog aggression depends on the temporal parameters of the rival, but also on how territory-holders compare their own calls to those of challengers in terms of their frequency, and therefore demonstrate that frogs pay attention to different aspects of signals during aggressive decision-making.

## Introduction

When animals are challenged by a rival, they must decide how to act based on the signals and cues they perceive from contenders. Adequate rival assessment is particularly important among animals that stablish and defend territories, as losing control over a territory can have important consequences for the survival and reproduction of the loser and therefore of the winner too (Ord 2021).

Behavioural studies have identified multiple factors influencing the level of aggression displayed by animals when challenged by rivals. These factors include the familiarity between the interacting individuals (i.e., the dear enemy effect, Temeles 1994), the prior experience with the territory (i.e., the resident effect, Yang et al., 2020), the hormonal state of interactants (i.e., the challenge hypothesis, Wingfield 2017), and demographic variables like sex-ratio, density of individuals or the presence polymorphisms in the population (e.g., Spence & Smith 2005, Yang et al., 2018), among other variables. All these factors are largely variable across species, populations, individuals, and even within individuals depending on the time of the day or the immediate social environment (e.g., Gardner & Graves 2005, Montroy et al., 2016). This multi-layered variability is at the centre of the difficulty to accurately predict how animals will behave during agonistic encounters in a generalizable way. Indeed, it has been recently argued that rival assessment strategies are not a species- or population-specific attribute, but rather an individual-level characteristic (Chapin et al., 2019), and therefore deserve to be studied in a case-by-case basis.

Another factor making it difficult to generalize how animals behave during aggressive interactions is that different animals rely on different sensory modalities to evaluate their rivals. Direct assessment is possible if the two contenders can, for example, see or touch each other, and therefore estimate the fighting capacity or body size of one another. However, in many cases the detection of a rival occurs through indirect signals or cues, like scents or sounds. In these cases, an initial assessment of a rival must be based on what an animal can indirectly perceive. Consequently, a lot of research has been devoted to establishing links between the physical attributes of animals (i.e., size or physiological state) and the characteristics of their signals (i.e., the frequency and loudness of their calls, or the pheromones in their scents) (reviewed in Searcy & Nowicki 2005). Many times, signals are an accurate reflection of the condition of the emitter, but this may not always the case, even when animals have direct access to perceiving their rivals (e.g., the size of a regenerated claw is not indicative of its strength in a crab, Bywater et al., 2015). When the link between the structure of signals and the attributes of individuals producing them is weak or absent they are unreliable indicators, and their influence on aggressive decision-making is unclear.

In the acoustic domain, one important indicator of the attributes of a signaller is the vocal performance, at least among birds. Across and within bird species, the trill rate and the frequency bandwidth of the vocalizations cannot be simultaneously maximized because of motor constraints on vocal production (e.g., Podos 1997, 2001). In other words, some combinations of these vocal traits are challenging or even impossible for birds to produce, and individuals calling closer to the empirical limit are considered high performers. Behavioural experiments have shown that birds respond differently to playbacks of calls with different performance levels in both territorial and reproductive contexts (e.g., Ballentine et al., 2004, de Kort et al., 2008), providing support to the idea that vocal performance acts as a reliable indicator of signaller quality (Podos 2017). Since its original formulation, the performance hypothesis has been expanded beyond birds to include other highly vocal animals, like mammals (e.g., Clink et al., 2018, Sun et al., 2022). Among frogs, however, vocal constraints on performance have been less studied. To the best of our knowledge, the only available example corresponds to the grey tree frog (*Hyla versicolor*), where a vocal trade-off equivalent to the one reported for birds has been found between call rate and call duration (Reichert & Gerhardt 2012).

Frogs are an excellent group for studying aggression. Anuran contests occur mostly between males during the breeding season (but see Rojas & Pašukonis 2019 for an example of female-female aggression), and typically take the form of vocal contests. These interactions can remain purely vocal, and some species have a distinctive aggressive call that they produce during escalated conflicts. In some cases, encounters escalate to physical combat, and can result in injury when species possess weapons like keratinized spines or fangs (Tsuji & Matsui 2002, Hudson et al., 2010). Even when contestants cannot harm each other, losing control over a territory can have important effects on the reproductive output of the territory holder if, for example, eggs are laid in the territory and their development depends on parental care (e.g., Quiguango-Ubillús & Coloma 2008), or there are favourable calling sites in the territory (e.g., Rodríguez et al., 2020).

Here we studied aggressive decision-making in a territorial poison-dart frog, the rainforest rocket frog (*Silverstoneia flotator*). Male *S. flotator* are small (around 15 mm snout-vent length) and call mostly from the top of leaves within their territories on the forest floor of Central America (Muñoz & Halfwerk 2022, **Fig. 1a**). Like many other species in the poison-dart frog family (Dendrobatidae) (Summers 2000, Pröhl 2005), male *S. flotator* hold and defend territories. This species lacks a distinctive aggressive call, and therefore advertisement calls are involved in both male-male and male-female interactions. Detecting the advertisement vocalization of another individual is enough to trigger the aggressive responses of territorial males. Their advertisement vocalizations are long and lasts several seconds. At the beginning of the call, notes are produced at a slow and irregular pace, but after this introductory phase they become more regular and the inter-note intervals shorter. All the males increase the note repetition rate as their calls progresses until reaching inter-note intervals of around 0.11 – 0.15 seconds, but never going below 0.1 seconds (**Fig. 1b,c, Supp. Fig. 1**, see the **Methods** section for a more detailed description of calls). Each note corresponds to an exhalation, and this threshold is probably the result of a physiological limit on the speed at which males can produce the notes (i.e., the rate at which they can contract their trunk muscles). Plotting the note repetition rate (notes/sec) of the fast portion of the calls against the duration of the complete call (introduction + fast sections) reveals the presence of an upper limit to their combination, and indicates that fast repetition rates are incompatible with very long calls (**Fig. 1d, Supp. Fig. 2**), similar to what has been reported for grey tree frogs (Reichert & Gerhardt 2012). Interestingly, the songs of the congeneric *S. nubicola* and some species from the sister genus, *Epipedobates*, also consist of the fast repetition of short notes, but these are emitted with inter-note intervals shorter than 0.10 seconds limit present in *S. flotator,* and their calls tend to be shorter (Ibáñez & Smith 1995, Scherges & Rödder 2017, **Supp. Fig. 3**), suggesting an inter-specific expression of the same performance trade-off.

**Figure 1:**
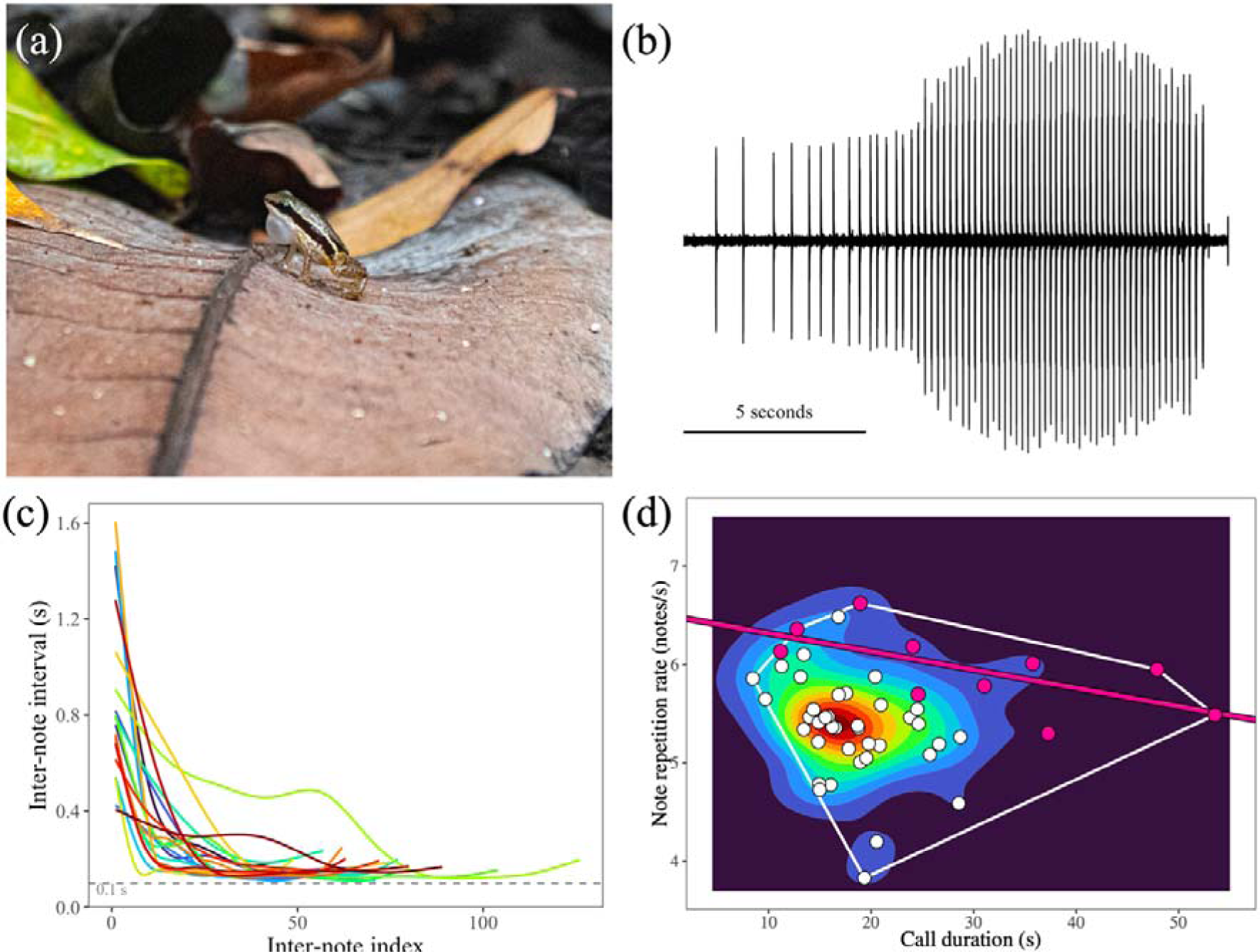
(a) Calling *S. flotator* male. (b) Example of the oscillogram of an advertisement call. (c) Inter-note interval as a function of the inter-note index (i.e., the ordination of all the inter-note intervals from the first to the last of a call) for 20 different rocket frogs. For visualization purposes the inter-note intervals are smoothed with a generalized additive model function (see **Supp. Fig. 1** to see the raw data). The grey horizontal line depicts the 0.1 seconds threshold. (d) Evidence for a performance limit on the call duration and note repetition rate of rocket frog calls (N = 50 individuals). The pink line depicts the upper-bound regression (slope = −0.02, P = 0.0396, r^2^ = 0.43), and the pink dots correspond to the data used to compute it. Further details about how the figure in panel (d) was obtained can be found in the **Supp. Fig. 2**.

Manipulating stimuli beyond the natural variation has been important to our understanding of the factors responsible for the evolution of signals, especially when these are manipulated in relation to the phylogenetic history of a group (Ryan & Cummings 2013), while testing behavioural responses to stimuli modified along one axis of the vocal performance space has contributed to our understanding of the behavioural significance of vocal trade-offs (e.g., de Kort et al., 2008). Based on the presence of an apparent performance limit on the calls of rocket frogs, and in consideration of the vocalizations of other closely related species, here we tested territorial aggression towards synthetic calls having faster and slower note repetition rates relative to the 0.1 seconds boundary, and that, therefore, simulated low- and exceptionally high-performance challengers.

## Materials and Methods

### Study site and call description

The experiments were performed during June and July of 2023 at Barro Colorado Island (BCI), Panamá, at the field station belonging to the Smithsonian Tropical Research Institute.

Below we provide a brief description (mean ± SD) of the rocket frog advertisement call based on the values measured from the vocalizations of 20 individuals recorded on 2019 and 2022 at BCI. The note is the basic unit of rocket frogs call, and have a duration of 0.024 seconds (± 0.004 s) and a peak frequency of 6.52 kHz (± 0.21 kHz). The notes are frequency modulated, and the frequency at the beginning and end of the notes are 6.23 kHz (± 0.189 kHz) and 6.75 kHz (± 0.21 kHz), respectively (measured from the frequency bandwidth at - 12 dB from the peak). Notes are arranged in complete calls that have a total duration of 19.46 seconds (± 8.20 s). The total duration of calls can be split in two section. An initial introductory section with a duration of 11.17 seconds (± 7.62 s), during which notes are produced at a slow and irregular rate (inter-note interval = 0.49 ± 0.20 s), followed by a final section with a duration of 8.29 seconds (± 1.76 s) and during which notes are repeated at a faster and more regular rate (inter-note interval = 0.16 ± 0.03 s). The introductory and final sections of calls form a continuum, but see **Supp. Fig. 2** for further details about the method used to stablish the transition between the introductory and final sections of calls. The amplitude of the notes at the final section of the calls is 88.5 dB SPL RMS (ref. 20 µPa) at 48.5 cm (N = 15 individuals), as computed from the recordings obtained from a microphone calibrated with the 114 dB SPL RMS 1-kHz pure tone delivered from a pistonphone (G.R.A.S. Sound calibrator type 42AB).

### Synthetic stimuli creation

We used Audacity (version 3.1.0, Audacity Team 2021) to create a synthetic rocket frog note based on the calls of the twenty individuals described above. The synthetic note had a duration of 0.026 seconds with a linear fade in and fade out of half the length of the note, and peak frequency of 6.46 kHz (**Fig. 2a**). The frequency of the note increased linearly from the beginning to the end from 6.1 to 6.8 kHz. The note was the basic unit used to create both the fast and slow synthetic calls. The two synthetic calls had a total duration of 17.9 seconds, with an introductory section corresponding to the first 6.3 seconds (**Fig. 2b**). The introductory section was identical between fast and slow stimuli, and the notes in this section had a linear rise in amplitude accompanied by a linear decrease in the inter-note intervals (i.e., by the end of the introductory section notes were of higher amplitude and repeated faster). We chose to use an introductory section shorter than the natural average (mean = 11.17 seconds) based on the fact that this section is largely variable across individuals, and on our aim to test the responses to the final portion of the synthetic calls instead of the introductory section. In natural calls, increases in the amplitude of the notes are accompanied by an increase in peak frequency (**Supp. Fig. 3**), however, we chose to keep the peak frequency of the synthetic note fixed for simplicity. The fast and slow stimuli differed only in the inter-note intervals of the final section of the calls. The slow call had inter-note intervals of 0.25 seconds, while the fast call had inter-note intervals of 0.05 seconds (**Fig. 2b, Supp. Fig. 5**). In terms of standard deviations (SD), the fast stimulus was +3.5 SD higher than natural calls (mean = 0.156 seconds, SD = 0.027), and the slow stimulus was −4.0 SD lower. This final section of the calls had a duration of 11.6 seconds. Overall, we are confident our synthetic calls were adequate to test aggressive responses towards stimuli largely deviating from the population mean in the temporal domain, while remaining close to the population average in the spectral domain (**Supp. Fig. 6**). During the playbacks, both synthetic calls were emitted with an inter-call interval (i.e., the silence gap between two consecutive calls) of 10 seconds. Both synthetic stimuli were calibrated so the notes at the final section of the calls reached an average amplitude of 90.6 dB SPL RMS (re. 20 μPa) at a distance of 60 cm on the forest floor.

**Figure 2:**
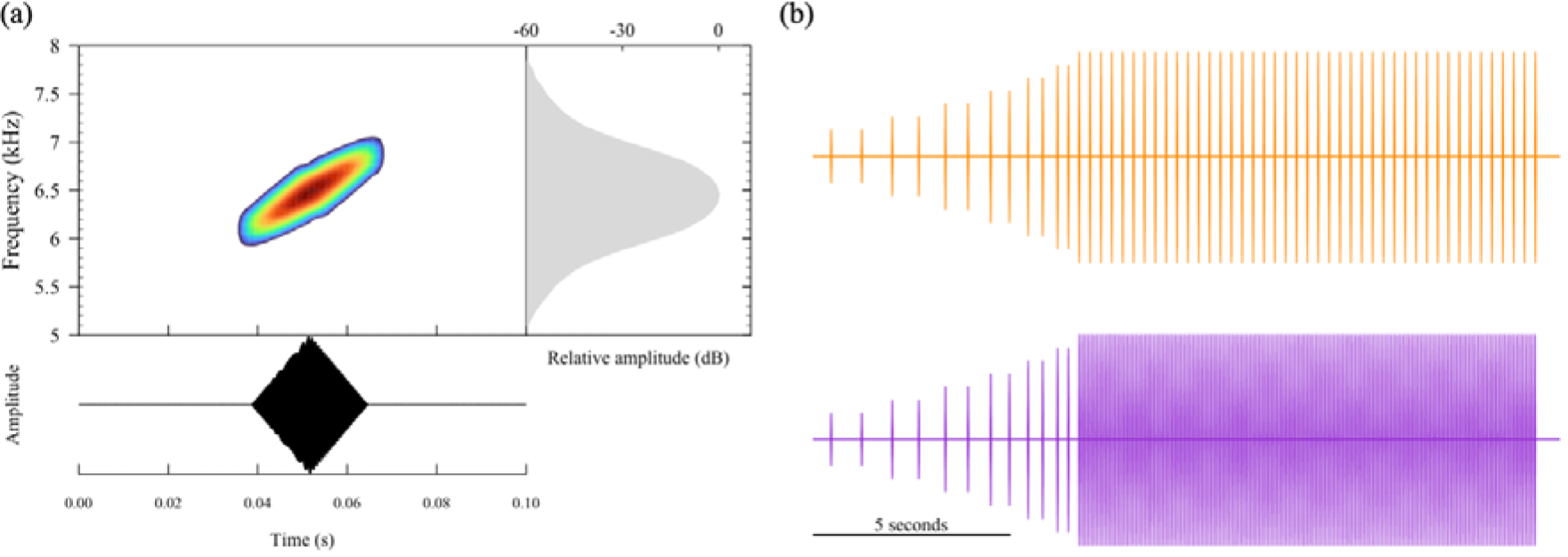
(a) Spectrogram (colourful, top left) power spectrum (grey, top right) and oscillogram (black, bottom left) of the synthetic rocket frog note. (b) Slow (top, orange) and fast synthetic calls (bottom, violet).

### Behavioural experiments

Actively calling males were located in the field during the peak activity periods of the species: in the morning after sunrise (7:00 to 12:00 AM) and the afternoon before sunset (15:00 to 17:30 PM). The territories of each male tested (N = 45) were identified with flagging tape, and we used these markings to avoid testing the same male twice. For each male we recorded their basal vocal activity for 5 minutes in audio (44.1 kHz, 16 bits, Sennheiser ME66 with K6 power module connected to a TASCAM DR-40 digital recorder) and video (Panasonic HC-V785). Males were tested only if they called at least once during this 5-minutes period. After recording their basal activity, each male was exposed to either the slow or the fast stimulus. Stimuli were broadcast from a small bluetooth speaker (3 cm diameter, BoomPod Zero connected to MECHEN M3-EU mp3 player) placed facing the focal male at a distance of between 36 to 85 cm (mean = 64.9 cm). The speaker was sitting on the centre of a 19 cm diameter cork platform that defined the response threshold. Males were exposed to the playback for a maximum of 10 minutes, or until an aggressive response was observed. A response was scored as aggressive once the male jumped on top of the cork platform (**Supp. Fig. 7**). The experiment was concluded earlier if the frogs moved away and disappeared from the sight of the experimenter for longer than 3 minutes. The vocal activity and behaviour of the males was recorded throughout the experiment. After finishing an experiment, we measured the distance between the speaker and the position of the frog at the onset of stimulation, and the temperature and humidity at the forest floor (Kestrel 5500). The frogs were not captured after the experiment. The starting distances between the speakers and the frogs were the same between the fast (mean ± SD. = 64.0 ± 8.90 cm) and the slow (mean ± SD = 65.8 ± 7.40 cm) acoustic treatments (two sample t-test: *t*_43_ = −-0.735, *P* = 0.467). Environmental conditions were also the same between both acoustic treatments (mean temperatures: 27.96 vs. 27.90 °C; mean humidity: 93.79 vs. 94.08 %, two sample t-tests: *P* > 0.70). We therefore rule out that different experimental conditions during stimuli presentation could have influenced our results.

Additionally, we recorded the spontaneous vocalizations of 14 males for 5 minutes. These males were different from the ones used for playback experiments. Unlike the males exposed to the playbacks, these males were captured and their body weight measured with a portable digital scale (Ohaus YA102, 0.01g precision). Individuals were released at the same place where they were captured after measuring them. These data were used to evaluate the relation between the weight and the acoustic traits of male calls.

### Ethical approval

All the experiments were licensed and approved by the Smithsonian Tropical Research Institute (IACUC SI-23011, approval date 24 February 2023).

### Data analysis

We scored the responses as either i) aggressive, if the frogs crossed the response threshold, or ii) non-aggressive, if the frogs stayed on the same place, approached the speaker but did not cross the response threshold, or moved away from the speaker. For those frogs that responded aggressively, we measured response latency (in seconds) as the time between stimulus onset and the moment they crossed the edge of the cork platform that defined the response threshold. One individual exposed to the slow call responded aggressively 11 seconds after the 10 minutes of playback while the stimulus was still being played. We kept this individual in our analysis to avoid reducing our sample size.

For each male we manually selected the last ten notes from three randomly chosen calls emitted during the basal activity period using RavenPro (version 1.6.5, Cornell Lab of Ornithology), except for two individuals with only one baseline call available (total of 1310 notes from 131 calls belonging to 45 individuals). These selections were used to measure the inter-note interval of male calls and the peak frequency of their notes (computed from the power spectrum, Hanning window, 512 samples, 90% overlap). We averaged both acoustic variables to a single value per individual for further analyses.

The majority of the males vocalized during stimulus presentation (41 out of 45 individuals). These calls were mostly emitted during the 10 seconds silence period between two consecutive synthetic calls. This silence gap was often too short for males to produce full calls and they would quickly stop calling when the stimulus began playing back again (i.e., frogs avoided overlapping with the stimulus, **Supp. Fig. 8**). Most of the males would not reach the final section typical of rocket frog calls, and therefore, we could not analyse the last ten consecutive notes of the calls as we did for the vocalizations produced during the basal activity period. Still, we selected 10 consecutive notes from one to three different calls produced just at the end of the 10-seconds silence gap between two synthetic calls (1000 notes from 100 calls belonging to 41 individuals). We could not measure inter-note intervals for these notes in a comparable way to the notes measured during the basal period, but we nevertheless measured their peak frequency, and used these data to evaluate if males modify their peak frequency in response to the playback.

We analysed the calls of the 14 males that were weighted in the same way as those recorded during the basal activity period of frogs exposed to the playbacks. In brief, we measured the inter-note intervals and peak frequencies of 10 to 30 notes from the end of one to three calls, and these data were averaged to a single value per individual.

### Statistical analyses

All the statistical analyses were done in R (version 4.3.1, R Core Team 2023), and figures were done using the libraries ‘*ggplot2*’ (version 3.4.2, Wickham 2016) and ‘*seewave’* (version 2.2.1, Sueur et al., 2018). We compared the proportion of frogs that responded aggressively towards the fast versus the slow stimulus using a Chi-square test. We compared the latency to respond between both acoustic treatments using the Wilcoxon signed rank test.

We evaluated if the probability of responding aggressively was related to the acoustic properties of calls produced during the basal activity period. For this we fitted a generalized linear model with the behavioural responses as a binary dependent variable (aggressive = 1, non-aggressive = 0) and a binomial distribution with a logit link function (i.e., a logistic regression). As predictors we included the acoustic treatment (categorical variable with two levels: fast or slow), the vocal attribute of the males (continuous variable: either inter-note interval or peak frequency), and their interaction as predictors. Our results suggested that males responded to the stimulus treatment based on the relative comparison of their own peak frequency against the peak frequency of the synthetic call (see *Results* section). Therefore, we transformed the absolute peak frequency of the males into two relative measures of peak frequency, a continuous and a categorical measure. We computed relative peak frequency as a continuous variable by subtracting the peak frequency of the stimulus (i.e., 6460 Hz) from the peak frequency of each male, and as a categorical variable by classifying males as calling either at lower or higher peak frequency than the stimulus (i.e., a factor with two levels). We fitted two additional logistic regressions identical to the one described above but including these relative measures of peak frequency as predictors instead of the absolute peak frequency of the individuals.

For those frogs that responded aggressively, we also tested whether the latencies to respond were related to their vocal attributes measured during the 5-min period of basal activity. For this we fitted linear regressions with the (log10-transformed) response latencies as dependent variable, and included the interaction between acoustic treatment and the vocal attributes of the males as predictors, similarly to the logistic regression described above. For the regression analyses of inter-note intervals (logistic and simple linear), we excluded two individuals that had outstandingly long intervals (< 0.20s), as these were clear outliers from the rest of the tested animals (**Supp. Fig. 9**). The results we report in the main text exclude those two individuals, but they were qualitatively the same as when these individuals were not excluded from the analyses (**Supp. Tables 1 and 2**).

To evaluate whether frogs modified the peak frequency of their call during playback exposure we fitted a linear-mixed effects models with the library ‘*lme4’* (version 1.1-34, Bates et al., 2015). We included peak frequency as a continuous response variable, the playback period (categorical variable with two levels: basal and playback), the relative peak frequency of males as a categorical variable (higher or lower than the stimulus), the stimulus treatment (categorical variable with two levels: fast or slow), and their triple interaction as fixed-effects. Individuals were included as a random effect to account for the repeated measures structure of the data. We computed *P*-values for the fixed-effects of this full model using the Satterthwaite’s method as implemented in the library ‘*lmerTest’* (version 3.1-3, Kuznetsova et al., 2017). This analysis showed no effect of stimulus treatment either by itself or in its interaction with the other fixed-effects (**Supp. Table 3**), and therefore we dropped it from the final model. For the final model we pooled together the data from males exposed to both acoustic treatments, and included the playback period (categorical variable with two levels: basal and playback), the relative peak frequency of males as a categorical variable (higher or lower than the stimulus), and their interaction as predictors. *Post hoc* comparisons were performed using the library ‘*emmeans’* (version 1.8.7, Lenth 2023).

Finally, for the 14 males that were recorded and weighted, we fitted two separate linear regressions to test if their peak frequencies and the inter-note intervals were related to their body weight. We visually inspected the residuals of all the linear models to check for deviations from the assumptions.

## Results

We testes 23 males with the fast stimulus, and 22 with the slow stimulus. Half of the frogs responded with aggression towards the fast stimulus (52%), while the majority of the males exposed to the slow stimulus tended to show aggressive responses (73%) (**Fig. 3a**). These two proportions were not significantly different from each other (χ^2^_1_ = 2.02, *P* = 0.155). Aggressive frogs had a 2.4-times shorter latency time to reach the speaker playing back the slow call (N = 16, median ± SD = 48.5 ± 151.9 seconds) than they reached the speaker playing back fast calls (N = 12, median ± SD = 118.0 ± 149.2 seconds, Wilcoxon rank sum test: *W* = 160, *P* = 0.003, **Fig. 3b**).

**Figure 3:**
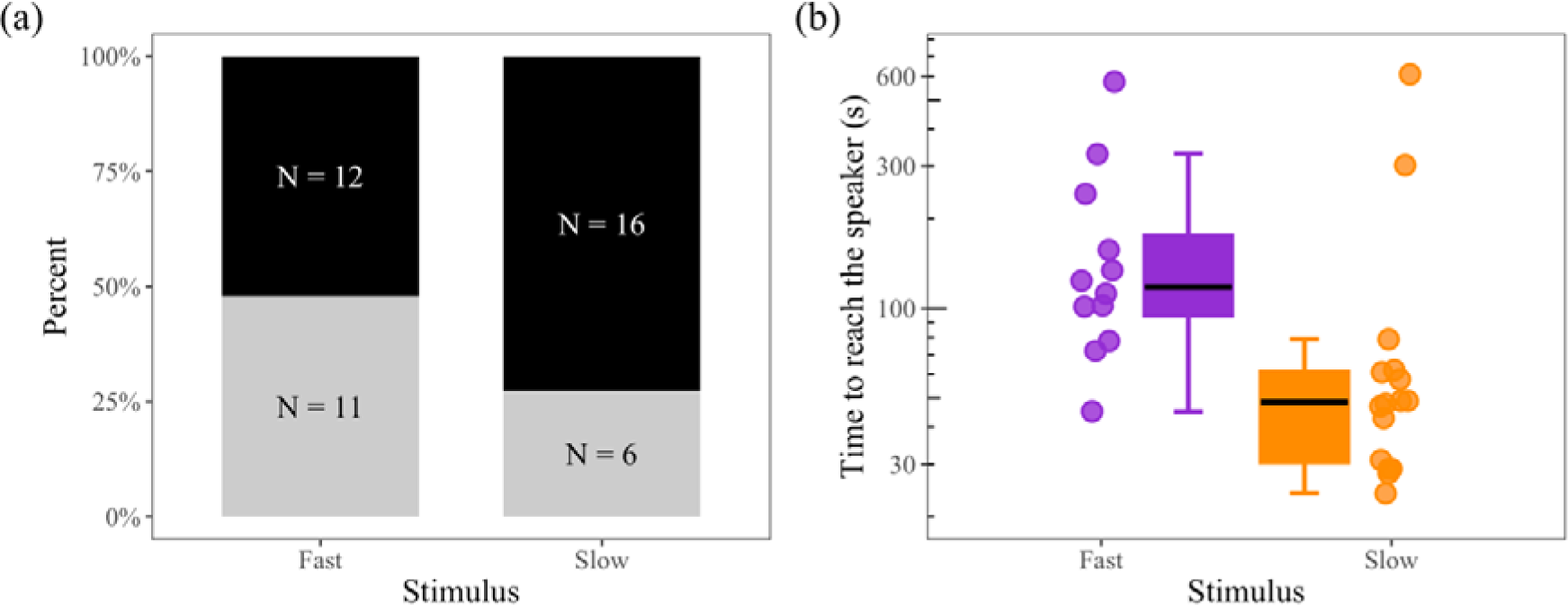
(a) Percent of aggressive (black) and non-aggressive (grey) responses given by males exposed to the fast and slow synthetic calls. (b) Response latencies (i.e., time to cross the response threshold) of males exposed to the fast (purple) and the slow (orange) stimulus. Note latencies are plotted on a logarithmic space.

Next, we evaluated whether the probability of aggression and the latency to respond were related to the call properties of the territory holder recorded during the 5-minutes basal activity period. We did not find an effect of the stimulus, the males’ inter-note interval, or their interaction, on the probability of responding aggressively (**Fig. 4a**, **Table 1**). Instead, we found that the probability of aggression depends on the interaction between the stimulus and the peak frequency of the territory holder (**Fig. 4b**, **Table 1**). Aggressive responses to the fast stimulus were more likely in males that called with lower peak frequencies than the synthetic note (i.e., 6.46 kHz), and less likely if they called with higher peak frequencies than the stimulus (**Fig. 4b**). The inverse occurred in response to the slow call: the probability of aggression was higher for males singing at high frequencies, and low for males singing at low frequencies (**Fig. 4b)**. For a male with a peak frequency of 7.0 kHz (i.e., a high-frequency calling male), the logistic model predicts a 11.1% probability of aggressive response towards the fast call, and a 95.3% probability of aggression towards the slow call. For a male with a peak frequency of 6.0 kHz (i.e., a low-frequency calling male), we predict an 88.1% probability of aggression towards the fast call, and a 25.5% probability of aggression towards the slow stimulus. We found the same significant interaction with stimulus when using relative peak frequency as continuous (*P* = 0.019) and categorical (*P* = 0.014) predictors in the logistic model (**Supp. Table 4**). Neither inter-note interval (**Fig. 4c**) nor peak frequency (**Fig. 4d**), or their interaction with stimulus, had a significant effect on the latency to respond (**Table 2**).

**Figure 4:**
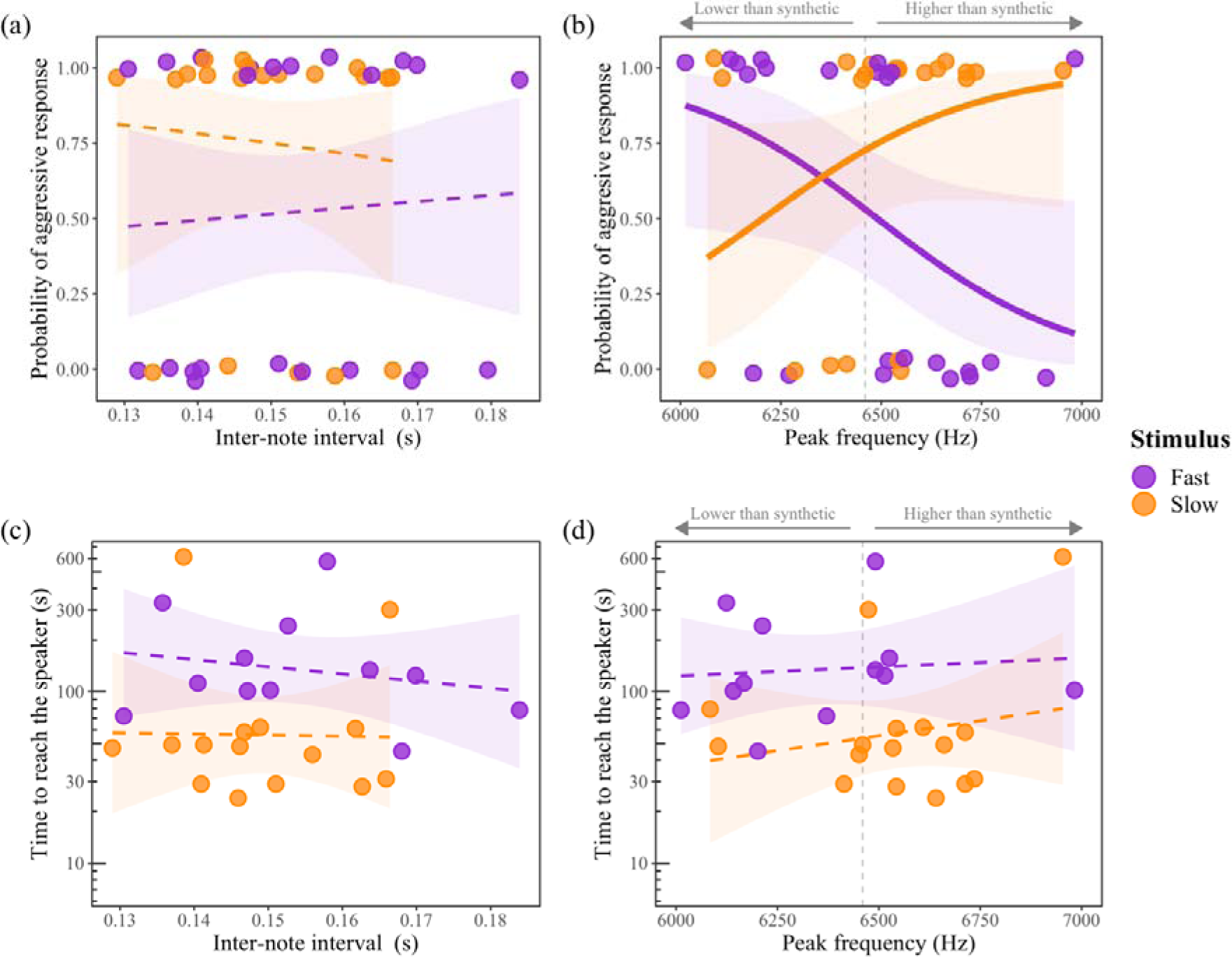
(Top row) Logistic regressions showing the probability of responding aggressively as a function of stimulus and the (a) inter-note interval and (b) peak frequency of the calls of the territory holder. Dashed regression lines in (a) indicate the lack of effect of inter-note interval, stimulus or their interaction on the probability of aggression. Points are slightly jittered vertically to improve visualization, but the response variable was binary. (Bottom row) linear regressions showing the lack of association between the times to reach the speaker and the (c) inter-note interval and (d) peak frequency of spontaneous male calls. Dashed regression lines in (c) and (d) indicate that the slopes were not different from 0. Note times are plotted on a logarithmic space. For reference, the vertical dashed grey line in (b) and (d) shows the peak frequency of the synthetic rocket frog note (6460 Hz).

**Table 1:** Results of logistic regressions fitted to evaluate whether the probability of responding aggressively depends on the vocal attributes of tested males (inter-note interval and peak frequency) and the stimulus.

| Vocal trait |  | Estimate | 95% CI | S.E. | z-value | P |
| --- | --- | --- | --- | --- | --- | --- |
| Inter-note interval | Intercept | -1.187 | [-9.936, 7.278] | 4.276 | -0.278 | 0.781 |
|  | Inter-note interval | 8.327 | [-46.701, 65.485] | 27.829 | 0.299 | 0.765 |
|  | Stimulus:Slow | 4.955 | [-11.155, 22.146] | 8.292 | 0.598 | 0.550 |
|  | Inter-note interval x Stimulus:Slow | -26.05 | [-138.040, 81.217] | 54.597 | -0.478 | 0.633 |
| Peak frequency | Intercept | 26.514 | [3.191, 57.252] | 13.312 | 1.992 | **0.046** |
|  | Peak frequency | -0.004 | [-0.009, -0.0005] | 0.002 | -1.988 | **0.047** |
|  | Stimulus:Slow | -50.382 | [-95.770, -12.098] | 20.890 | -2.412 | **0.016** |
|  | Peak frequency x Stimulus:Slow | 0.008 | [0.002, 0.015] | 0.003 | 2.450 | **0.014** |

S.E. = standard error. Significant factors are in bold.

**Table 2:** Results of the linear regressions fitted to evaluate if latencies to respond aggressively depends on the vocal attributes of tested males (inter-note interval or peak frequency) and stimulus.

| Vocal trait |  | Estimate | 95% CI | S.E. | t-value | P |
| --- | --- | --- | --- | --- | --- | --- |
| Inter-note interval | Intercept | 2.772 | [0.497, 5.047] | 1.100 | 2.52 | **0.019** |
|  | Inter-note interval | -4.180 | [-18.894, 10.533] | 7.113 | -0.588 | 0.562 |
|  | Stimulus:Slow | -0.923 | [-4.431, 2.584] | 1.696 | -0.545 | 0.591 |
|  | Inter-note interval x Stimulus:Slow | 3.504 | [-19.621, 26.628] | 11.178 | 0.313 | 0.757 |
| Peak frequency | Intercept | 1.466 | [-3.813, 6.746] | 2.558 | 0.532 | 0.572 |
|  | Peak frequency | 0.0001 | [-0.0007, 0.0009] | 0.0004 | 0.259 | 0.798 |
|  | Stimulus:Slow | -1.988 | [-9.697, 5.720] | 3.735 | -0.532 | 0.599 |
|  | Peak frequency x Stimulus:Slow | 0.0002 | [-0.0009, 0.0014] | 0.0006 | 0.422 | 0.677 |

S.E. = standard error. Significant factors are in bold.

We found that males modified the peak frequency of their calls in response to the playback depending on whether they sang at higher or lower frequencies than the stimulus, as revealed by a significant interaction between playback period (basal and playback) and the relative frequency of male calls (higher or lower than synesthetic) (**Fig. 5**, **Table 3**). Males calling at lower frequencies than the stimulus increased their peak frequency in response to the playback (mean basal = 6258 Hz, mean playback = 6419 Hz, *post hoc* test: *t*-ratio_39_ = - 4.58, *P* < 0.001). In contrast, males that called at higher peak frequencies than the synthetic did not modify their calls in response to the stimulus (mean basal = 6630 Hz, mean playback = 6619 Hz, *post hoc* test: *t*-ratio_39_ = 0.38, *P* = 0.710).

**Figure 5:**
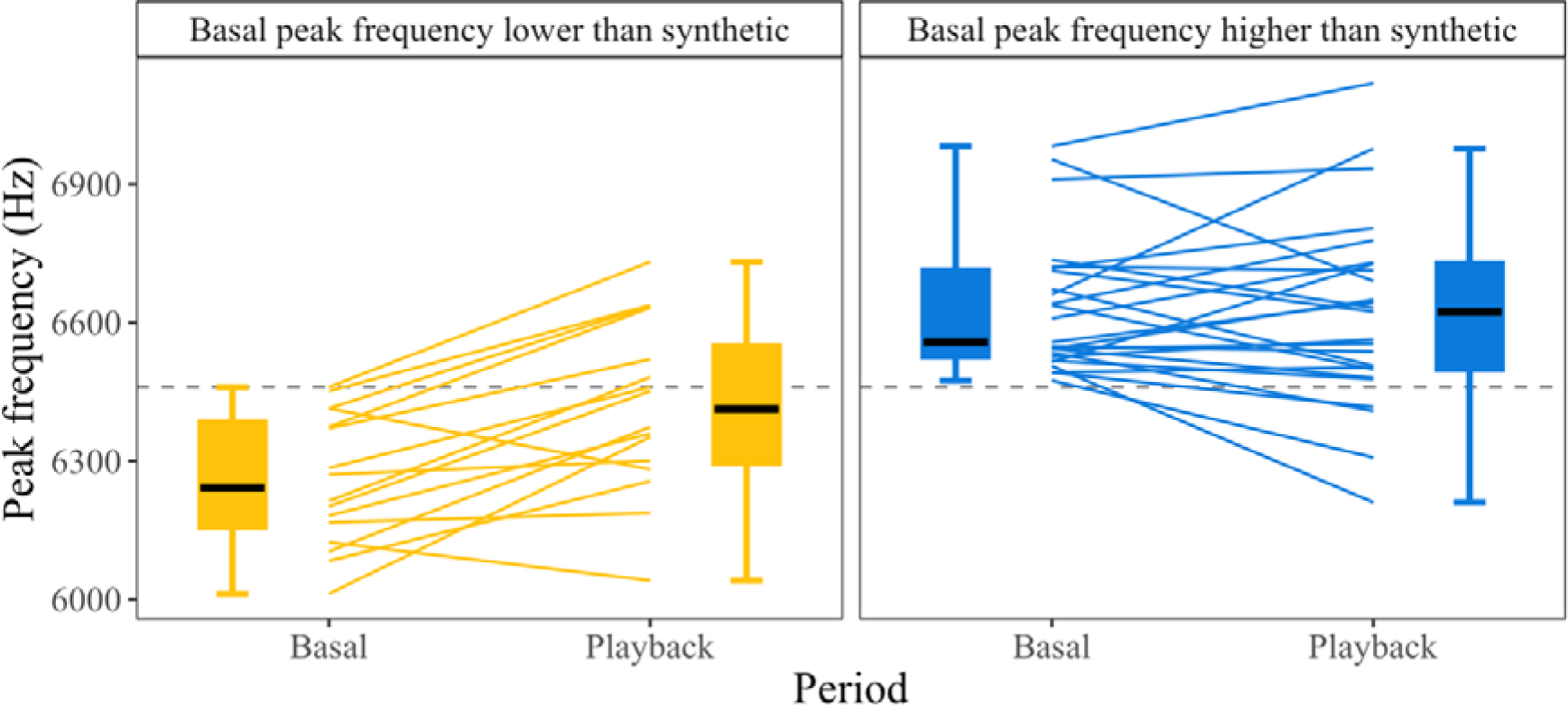
Comparison between the peak frequencies of calls emitted during the basal activity period and during the playback. Males that called with lower peak frequency than the synthetic call (6460 Hz, horizontal grey line) during the basal activity period are shown in yellow (left panel), and males that called at higher frequencies in blue (right panel). Lines connect the individual responses of each male.

**Table 3:** Results of the linear mixed-effects model fitted to evaluate differences in the peak frequency between calls emitted during the basal activity period and during the playback.

|  | Estimate | 95% CI | S.E. | d.f. | *t*-value | *P* |
| --- | --- | --- | --- | --- | --- | --- |
| Intercept | 6629.93 | [6562.412, 6697.447] | 34.72 | 53.79 | 190.976 | > **0.001** |
| Period:Playback | -10.57 | [-65.687, 44.556] | 28.16 | 39.00 | -0.375 | 0.710 |
| MaleFrequency:Lower | -372.02 | [-480.103, -263.941] | 55.57 | 53.79 | -6.694 | > **0.001** |
| Period:Playback x MaleFrequency:Lower | 171.62 | [83.379, 259.854] | 45.08 | 39.00 | 3.807 | > **0.001** |

S.E. = standard error, d.f. = degrees of freedom. Significant fixed-effects are in bold.

Neither peak frequency (linear regression: *F*_1,12_ = 1.633, *P* = 0.226, R^2^ = 0.1198) nor the inter-note interval of male calls (linear regression: *F*_1,12_ = 0.008, *P* = 0.929, R^2^ = 0.0007) were related to their body weight (**Fig. 6**).

**Figure 6:**
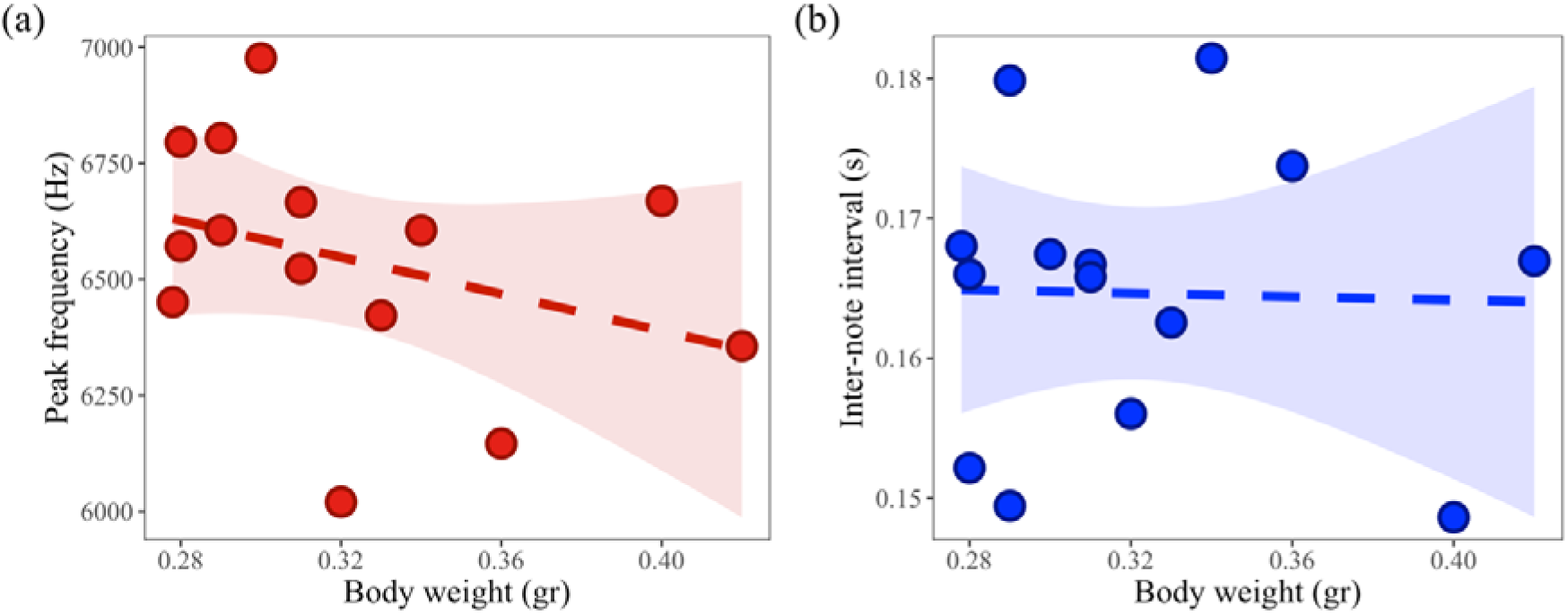
Allometry between body weight and peak frequency (a, red), body weight and inter-note interval (b, blue), and association between peak frequency and inter-note interval (c, orange). Dashed lines indicate that slopes are not different from zero.

## Discussion

When animals are challenged by rivals, they must decide how to act based on the signals and cues they can perceive from their challengers. Here we tested male aggression in response to synthetic calls designed to have faster and slower note repetition rates than those found in nature, effectively simulating a high- and a low-performance challenger. We found that the probability of aggression depended on the interaction between the stimulus treatment and the spectral properties the territorial males relative to the stimulus. The probability of aggression towards the fast call was negatively associated with the peak frequency of the territory-holder, meaning that low-frequency males were more likely to respond aggressively towards the fast call. The inverse was true for the slow vocalization, and aggression towards this stimulus was higher when males called at high peak frequency. We also found that males increase the peak frequency of their calls, but only if they called at lower frequency than the stimulus during the basal activity period. Body mass was not a reliable predictor of either the peak frequency, or the inter-note interval of male calls. Overall, our results show that temporal variables are important modulators of rocket frog aggression, and that males respond differently to variation in the performance of challengers depending on their own spectral (but not temporal) attributes.

Previous studies have shown that the rate of note or call production can be important for male-male interactions in frogs. For example, male *Allobates femoralis* have a wide recognition space in terms of inter-note intervals, and continue to respond with high levels of aggression towards calls with both higher and lower note repetition rates than the species average (Vélez et al., 2012). In contrast, male strawberry poison frogs (*Oophaga pumilio*) respond with elevated aggression towards average and low call rates, and low aggression towards high call rates (Protti-Sánchez et al., 2023), which is indicative of asymmetric responses towards fast and slow calls. The responses of frogs from the genus *Batrachyla* to exceptional variation in note or pulse rate are species-specific (Penna 1997, Penna et al., 1997, Penna et al., 2022). These studies indicate that broad recognition spaces in the temporal domain are a general feature of frog communication systems, although generalizing how individuals will respond to faster or slower rates is not straightforward. The fact that male rocket frogs responded to both of our stimuli agrees with such broad recognition spaces. In our experiment, males tended to be more likely to approach the slow stimulus (73 vs 52%, although the difference was not significant), and did it 2.4-times faster than to the fast call. We interpret these results as frogs perceiving the fast call as a more threatening rival, and therefore being less likely to risk a physical encounter. However, this is not the full picture, as our experiment also revealed that the probability of aggression is not only a function of the temporal variation of the stimulus, but also depends on the spectral properties of the territory holders.

Low-frequency calling rocket frogs showed higher aggression towards the fast (high-performance) synthetic call, while high-frequency calling individuals were more aggressive towards the slow (low-performance) call. Several studies have shown that male frogs respond with higher aggression towards high-frequency vocalization, and *vice versa*, with low aggression towards low-frequency ones (Davies & Halliday 1978, Arak 1983, Given 1987, Wagner 1989, Peignier et al., 2023, but see Bee 2002 and Morais et al., 2016 for exceptions). At first glance, our results seem to agree with these studies. However, one important component of these investigations was that the frequency content of male calls was negatively correlated with size or weight, and therefore aggressive responses to high-frequency rivals interpreted as an adaptive response toward smaller and less intimidating rivals. This was not the case of rocket frogs, where body weight and peak frequency were not correlated despite the wide range of inter-individual variation in peak frequencies, going from around 6.0 to 7.0 kHz. This makes it difficult to interpret our results in the same terms as the studies just mentioned. One possibility is that a heavier weight does not necessarily confer a competitive advantage. If body weight is not an important determinant of contest resolution, like it has been shown in *Hyla versicolor* (Reichert & Gerhardt 2011) and *Pithecopus nordestinus* (Brasileiro et al., 2020), then acoustic cues to body size would be irrelevant for aggressive decision-making, potentially resulting in the decoupled evolution of frequency and body size in the long term.

Another possibility is that peak frequency is associated to another vocal variable we did not measure, like amplitude. Within a rocket frog call, the amplitude and the peak frequency of the notes are positively associated (**Supp. Fig. 3**), and therefore, a higher-frequency challenger may be associated to a louder individual. This coupling between amplitude and frequency is present in birds (Zollinger et al., 2012), and in frogs is supported by excised larynx experiments showing that at higher pulmonary pressure, both the amplitude and the frequency content of the vocalization increases in a correlated fashion (Gridi-Papp 2014). Therefore, frogs can increase both the amplitude and frequency of their calls by pushing air out of their lungs more strongly.

Whether or not pushing air more strongly, and therefore calling louder and at higher frequencies, is indicative of the motivational state of an individual is unclear, but not an unlikely possibility. In our experiments, the microphone was not calibrated and the frogs were freely moving around the setup, making it difficult for us to reliably measure the amplitude of their calls. Testing responses to stimuli differing in peak frequency and presented at different amplitudes can help dissect the contribution of these two variables on rocket frog aggression in the future.

The fact that males were capable of modifying the frequency of their vocalizations also complicates the usefulness of peak frequency for rival assessment. Males from different frog species modify the spectral content of their calls during interactions with other males. The majority of these studies have reported that males either decrease the frequency of their calls in response to other males (Howard & Young 1998, Wagner 1989, Grafe 1995, Bee et al., 2000, Bee & Bowling 2002, Meuche et al., 2012, Reichert & Gerhardt 2013), or increase or decrease their frequency to match that of their opponent (Lopez et al., 1988). Most of our rocket frog males that called at lower frequencies than the stimulus increased their peak frequency during the playback (14 out of 16 individuals) by an average of 161.1 ± 134.9 Hz. In contrast, around the same number of males that called at higher frequencies than the stimulus decreased (13 out of 25 individuals) or increased (12 out of 25 individuals) their peak frequency during the playback, and the average change was −10.6 ± 144.4 Hz. This pattern of response is akin to the frequency matching described for white-lipped frogs (*Leptodactylus albilabris*) by Lopez et al., (1988), although it could be considered a form of partial matching because high-frequency males did not lower their peak frequencies to match the stimulus. In light of the association between frequency and amplitude mentioned above, and its potential link with motivation state, it is possible that the males spontaneously calling at low-frequencies were not singing at their full capacity, but switched to louder and higher frequencies calls when challenged. Males calling at higher-frequencies may have been at or near their full capacity, and therefore possibly could not substantially modify their calls.

Since the foundational study of Podos (1997), vocal performance studies on birds have flourished. Just like birds, frogs rely on vocal communication for important aspects of their lives, and their calls may be subject to motor constraints equivalent to those demonstrated for their avian counterparts. Yet, with the exception of Reichert & Gerhardt (2012), these ideas have not been widely transferred to the study of frog vocalizations. Here we found evidence in support of a trade-off between call duration and note repetition rate in rocket frogs, which is in line with the findings of Reichert & Gerhardt (2012). By manipulating note repetition rate while holding call duration constant we introduced a confounding variable in our experiments as the stimuli we used differed also in their total number of notes. We could have reduced the duration of the fast call to match the number of notes present in the slow call, but this would have resulted in call duration becoming a confounding variable instead of total note number. On top of that, rocket frogs approach (simulated) rivals during stimulus presentation because they need the sound to orient towards the intruder, and therefore testing synthetic call of different duration but same number of notes would have impacted our measures of response latencies. Different ways of synthetizing stimuli unavoidably introduce different confounding variables. Experiments testing multiple different combinations of these variables will be necessary to formally test whether frogs, for example, pay attention or not to the total number of notes in the calls of challengers. This is, nevertheless, to our knowledge the first time that the aggressive behaviour of rocket frogs is studied, and the second time that performance limits have been demonstrated among frogs. Other studies should continue to evaluate the presence of these vocal production constraints and their consequences for aggressive and reproductive interactions within species, but also at the inter-specific level. The frogs from the genus *Silverstoneia* and their sister genus, *Epipedobates,* stand as good candidates for pursuing such a line of research.

## Conclusion

Here we simulated rivals with high and low vocal performances by manipulating the note repetition rate of synthetic rocket frog calls, and tested male aggression in response to these stimuli. We found that males have a wide recognition space in the temporal domain as shown by them responding to both of our stimuli. The probability of aggression to each acoustic treatment depended on whether males were on the higher or lower ends of peak frequency variation in the population, indicating frequency is an important feature in male-male interactions. Challenged males increased the peak frequency of their calls only if they sang at lower peak frequencies than the stimulus, but not if they did it at higher values. Call frequency and amplitude are mechanistically coupled in frogs, and are likely to reflect the motivational state of males. We argue that males calling at low-frequencies were not at their maximum calling capacity, which they increased in response to the playback, especially if the rival corresponded to a high-performance individual. If for any reason, an individual with an exceptionally high note repetition rate were to appear in the population, similar to the rates present in *Silverstoneia* or *Epipedobates* species, it would experience increased aggression from the low-frequency calling males, and these males would likely increase the frequency of their calls when challenged by this individual. Whether females would prefer such a male is unknown, but it is a possibility that deserves further testing.

## Author contributions

MM conceived the study with input from WH. MM and NC performed the experiments. MM analysed the data and wrote the manuscript with input from WH.

## Acknowledgements

We are thankful to the staff of the Smithsonian Tropical Research Institute on Barro Colorado Island for their fundamental support during data collection, and to Rachel Page, Roberto Ibáñez and William Wcislo for insightful conversations about frog behaviour. We are grateful to Mario Penna and Jacintha Ellers for providing valuable comments on early versions of the manuscript. MIM is thankful to the Cornell Lab of Ornithology for providing a free subscription to RavenPro. MIM is supported by Becas Chile 2018—CONICYT scholarship (number: 72190501), and the Dobberke Foundation for the Study of Behaviour (KNAW).

## References

Arak, A. (1983). Sexual selection by male–male competition in natterjack toad choruses. Nature, 306(5940), 261–262.

Audacity Team (2021). Audacity: Free Audio Editor and Recorder [Computer program]. Version 3.1.0. http://audacityteam.org.

Ballentine, B., Hyman, J., & Nowicki, S. (2004). Vocal performance influences female response to male bird song: an experimental test. Behavioral Ecology, 15(1), 163–168.

Bates, D., Mächler, M., Bolker, B., & Walker, S. (2015). Fitting Linear Mixed-Effects Models Using lme4. Journal of Statistical Software, 67, 1–48.

Bee, M. A. (2002). Territorial male bullfrogs (*Rana catesbeiana*) do not assess fighting ability based on size-related variation in acoustic signals. Behavioral Ecology, 13(1), 109–124.

Bee, M. A., & Bowling, A. C. (2002). Socially mediated pitch alteration by territorial male bullfrogs, *Rana catesbeiana*. Journal of Herpetology, 36(1), 140–143.

Bee, M. A., Perrill, S. A., & Owen, P. C. (2000). Male green frogs lower the pitch of acoustic signals in defense of territories: a possible dishonest signal of size?. Behavioral Ecology, 11(2), 169–177.

Brasileiro, A. C., Lima-Araujo, F., Passos, D. C., & Cascon, P. (2020). Are good fighters also good singers? The relationship between acoustic traits and fight success in the treefrog *Pithecopus nordestinus* (Phyllomedusidae). Acta Ethologica, 23(2), 51–60.

Bywater, C. L., Seebacher, F., & Wilson, R. S. (2015). Building a dishonest signal: the functional basis of unreliable signals of strength in males of the two-toned fiddler crab, *Uca vomeris*. Journal of Experimental Biology, 218(19), 3077–3082.

Chapin, K. J., Peixoto, P. E. C., & Briffa, M. (2019). Further mismeasures of animal contests: a new framework for assessment strategies. Behavioral Ecology, 30(5), 1177–1185.

Clink, D. J., Charif, R. A., Crofoot, M. C., & Marshall, A. J. (2018). Evidence for vocal performance constraints in a female nonhuman primate. Animal Behaviour, 141, 85–94.

Davies, N. B., & Halliday, T. R. (1978). Deep croaks and fighting assessment in toads *Bufo bufo*. Nature, 274(5672), 683–685.

de Kort, S. R., Eldermire, E. R., Cramer, E. R., & Vehrencamp, S. L. (2009). The deterrent effect of bird song in territory defense. Behavioral Ecology, 20(1), 200–206.

Gardner, E. A., & Graves, B. M. (2005). Responses of resident male *Dendrobates pumilio* to territory intruders. Journal of Herpetology, 248–253.

Given, M. F. (1987). Vocalizations and acoustic interactions of the carpenter frog, *Rana virgatipes*. Herpetologica, 467–481.

Grafe, T. U. (1995). Graded aggressive calls in the African painted reed frog *Hyperolius marmoratus* (Hyperoliidae). Ethology, 101(1), 67–81.

Gridi-Papp, M. (2014). Is the frequency content of the calls in North American treefrogs limited by their larynges?. International Journal of Evolutionary Biology, 2014.

H. Wickham. ggplot2: Elegant Graphics for Data Analysis. Springer-Verlag New York, 2016.

Howard, R. D., & Young, J. R. (1998). Individual variation in male vocal traits and female mating preferences in *Bufo americanus*. Animal Behaviour, 55(5), 1165–1179.

Hudson, C. M., He, X., & Fu, J. (2011). Keratinized nuptial spines are used for male combat in the Emei moustache toad (*Leptobrachium boringii*). Asian Herpetological Research, 2(3), 142–148.

Ibáñez, R., & Smith, E. M. (1995). Systematic status of *Colostethus flotator* and *C. nubicola* (Anura: Dendrobatidae) in Panama. Copeia, 446–456.

K. Lisa Yang Center for Conservation Bioacoustics at the Cornell Lab of Ornithology. (2023). Raven Pro: Interactive Sound Analysis Software (Version 1.6.5). Ithaca, NY: The Cornell Lab of Ornithology. Available from https://ravensoundsoftware.com.

Kuznetsova A., Brockhoff P.B., & Christensen R.H.B. (2017). lmerTest Package: Tests in Linear Mixed Effects Models. Journal of Statistical Software, 82(13), 1–26.

Lenth R (2023). emmeans: Estimated Marginal Means, aka Least-Squares Means R package version 1.8.7

Lopez, P. T., Narins, P. M., Lewis, E. R., & Moore, S. W. (1988). Acoustically induced call modification in the white-lipped frog, *Leptodactylus albilabris*. Animal Behaviour, 36(5), 1295–1308.

Meuche, I., Linsenmair, K. E., & Pröhl, H. (2012). Intrasexual competition, territoriality and acoustic communication in male strawberry poison frogs (*Oophaga pumilio*). Behavioral Ecology and Sociobiology, 66, 613–621.

Montroy, K., Loranger, M. J., & Bertram, S. M. (2016). Male crickets adjust their aggressive behavior when a female is present. Behavioural Processes, 124, 108–114.

Morais, A. R., Siqueira, M. N., Márquez, R., & Bastos, R. P. (2016). Males of *Hypsiboas goianus* (Anura; Hylidae) do not assess neighbor fighting ability through acoustic interactions. Acta Ethologica, 19, 43–50.

Muñoz, M. I., & Halfwerk, W. (2022). Amplification of frog calls by reflective leaf substrates: implications for terrestrial and arboreal species. Bioacoustics, 31(4), 490–503.

Ord, T. J. (2021). Costs of territoriality: a review of hypotheses, meta-analysis, and field study. Oecologia, 197(3), 615–631.

Peignier, M., Bégué, L., Ringler, M., Szabo, B., & Ringler, E. (2023). Regardless of personality, males show similar levels of plasticity in territory defense in a Neotropical poison frog. Scientific Reports, 13(1), 3435.

Penna, M. (1997). Selectivity of evoked vocal responses in the time domain by frogs of the genus *Batrachyla*. Journal of Herpetology, 202–217.

Penna, M., Feng, A. S., & Narins, P. M. (1997). Temporal selectivity of evoked vocal responses of *Batrachyla antartandica* (Amphibia: Leptodactylidae). Animal Behaviour, 54(4), 833–848.

Penna, M., Solís, R., & Moreno-Gómez, F. N. (2022). Diverse patterns of responsiveness to fine temporal features of acoustic signals in a temperate austral forest frog, *Batrachyla leptopus* (Batrachylidae). Bioacoustics, 31(2), 219–239.

Podos, J. (1997). A performance constraint on the evolution of trilled vocalizations in a songbird family (Passeriformes: Emberizidae). Evolution, 51(2), 537–551.

Podos, J. (2001). Correlated evolution of morphology and vocal signal structure in Darwin’s finches. Nature, 409(6817), 185–188.

Podos, J. (2017). Birdsong performance studies: reports of their death have been greatly exaggerated. Animal Behaviour, 125, e17–e24.

Pröhl, H. (2005). Territorial behavior in dendrobatid frogs. Journal of Herpetology, 39(3), 354–365.

Protti-Sánchez, F., García-Rodríguez, A., Barrantes, G., & Sandoval, L. (2023). Does variation in call rate affect the response of territorial males in the strawberry poison frog (*Oophaga pumilio*)?. Ichthyology & Herpetology, 111(2), 248–253.

Quiguango-Ubillús, A., & Coloma, L. A. (2008). Notes on behaviour, communication and reproduction in captive *Hyloxalus toachi* (Anura: Dendrobatidae), an Endangered Ecuadorian frog. International Zoo Yearbook, 42(1), 78–89.

R Core Team (2023). R: A Language and Environment for Statistical Computing. R Foundation for Statistical Computing, Vienna, Austria. https://www.R-project.org.

Reichert, M. S., & Gerhardt, H. C. (2011). The role of body size on the outcome, escalation and duration of contests in the grey treefrog, *Hyla versicolor*. Animal Behaviour, 82(6), 1357–1366.

Reichert, M. S., & Gerhardt, H. C. (2012). Trade-offs and upper limits to signal performance during close-range vocal competition in gray tree frogs *Hyla versicolor*. The American Naturalist, 180(4), 425–437.

Reichert, M. S., & Gerhardt, H. C. (2013). Gray tree frogs, *Hyla versicolor*, give lower-frequency aggressive calls in more escalated contests. Behavioral Ecology and Sociobiology, 67, 795–804.

Rodríguez, C., Amézquita, A., Ringler, M., Pašukonis, A., & Hödl, W. (2020). Calling amplitude flexibility and acoustic spacing in the territorial frog *Allobates femoralis*. Behavioral Ecology and Sociobiology, 74, 1–10.

Rojas, B., & Pašukonis, A. (2019). From habitat use to social behavior: natural history of a voiceless poison frog, *Dendrobates tinctorius*. PeerJ, 7, e7648.

Ryan, M. J., & Cummings, M. E. (2013). Perceptual biases and mate choice. Annual Review of Ecology, Evolution, and Systematics, 44, 437–459.

Scherges, A. M., & Rödder, D. (2017) The advertisement calls of *Epipedobates anthonyi* (Noble, 1921) and *Epipedobates tricolor* (Boulenger, 1899)(Anura: Dendrobatidae: Colostethinae): intra-and interspecific comparisons. Bonn Zoological Bulletin, 66(1), 73–84.

Searcy, W. A., & Nowicki, S. (2005). The Evolution of Animal Communication: Reliability and Deception in Signaling Systems: Reliability and Deception in Signaling Systems. Princeton University Press.

Spence, R., & Smith, C. (2005). Male territoriality mediates density and sex ratio effects on oviposition in the zebrafish, *Danio rerio*. Animal Behaviour, 69(6), 1317–1323.

Sueur, J., Aubin, T., & Simonis, C. (2008). Seewave, a free modular tool for sound analysis and synthesis. Bioacoustics, 18(2), 213–226.

Summers, K. (2000). Mating and aggressive behaviour in dendrobatid frogs from Corcovado National Park, Costa Rica: a comparative study. Behaviour, 137(1), 7–24.

Sun, C., Zhang, C., Lucas, J. R., Gu, H., Feng, J., & Jiang, T. (2022). Vocal performance reflects individual quality in male Great Himalayan leaf-nosed bats (*Hipposideros armiger*). Integrative Zoology, 17(5), 731–740.

Temeles, E. J. (1994). The role of neighbours in territorial systems: when are they ‘dear enemies’?. Animal Behaviour, 47(2), 339–350.

Tsuji, H., & Matsui, M. (2002). Male-male combat and head morphology in a fanged frog (*Rana kuhlii*) from Taiwan. Journal of Herpetology, 36(3), 520–526.

Vélez, A., Hödl, W., & Amézquita, A. (2012). Sound or silence: Call recognition in the temporal domain by the frog *Allobates femoralis*. Ethology, 118(4), 377–386.

Wagner, W. E. (1989). Fighting, assessment, and frequency alteration in Blanchard’s cricket frog. Behavioral Ecology and Sociobiology, 25, 429–436.

Wingfield, J. C. (2017). The challenge hypothesis: Where it began and relevance to humans. Hormones and Behavior, 92, 9–12.

Yang, Y., Dugas, M. B., Sudekum, H. J., Murphy, S. N., & Richards-Zawacki, C. L. (2018). Male–male aggression is unlikely to stabilize a poison frog polymorphism. Journal of Evolutionary Biology, 31(3), 457–468.

Yang, Y., Prémel, V., & Richards-Zawacki, C. L. (2020). Prior residence effect determines success of male–male territorial competition in a color polymorphic poison frog. Ethology, 126(12), 1131–1140.

Zollinger, S. A., Podos, J., Nemeth, E., Goller, F., & Brumm, H. (2012). On the relationship between, and measurement of, amplitude and frequency in birdsong. Animal Behaviour, 84(4), e1–e9.

