## Supplementary Figures and Tables for "Aggressive responses to rivals depend on the interaction between the vocal traits of territory-holder and mimicked intruders in a miniature tropical frog"

by

Matías I. Muñoz, Nicolás Camargo-Rodriguez & Wouter Halfwerk
Vrije Universiteit Amsterdam, The Netherlands

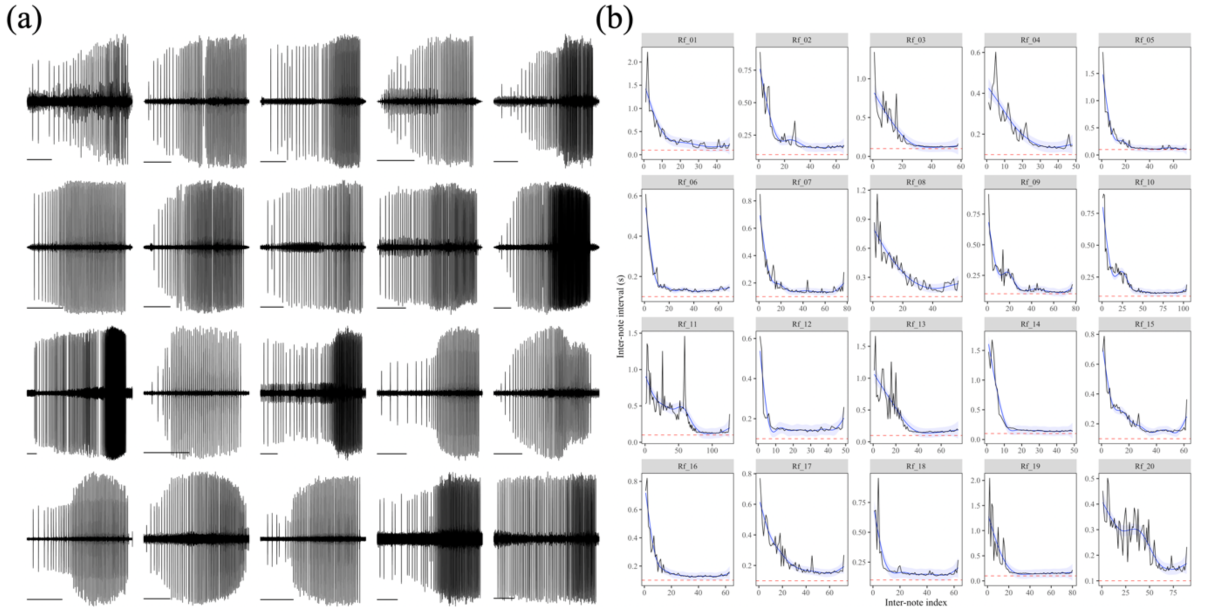


**Supplementary figure 1:** (a) Oscillogram of the advertisement calls of 20 rainforest rocket frog males. Scale bars = 5 seconds. (b) Inter-note interval versus inter-note index for the same 20 males as in (a). The black lines correspond to the raw inter-note intervals and the blue lines to the generalized additive model smoothing functions. The horizontal dashed red line corresponds to the 0.1 seconds inter-note interval threshold. These are the same calls shown in **Fig. 1C** of the main text. Note all panels in (b) have different scales.

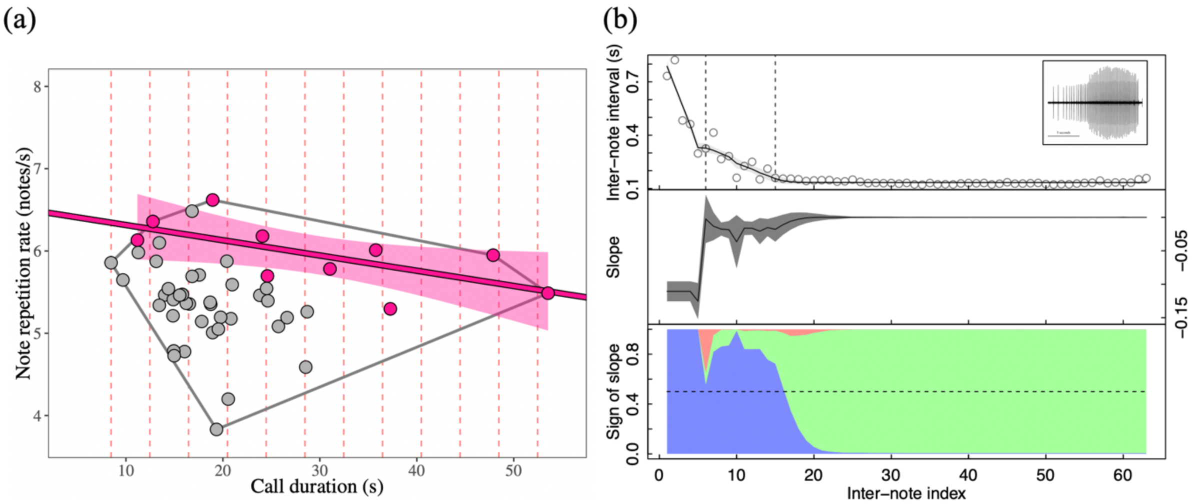


**Supplementary figure 2:** (a) Evidence for a performance constraint on rocket frog vocalizations. For each individual (N =50), we used the Bayesian change point detection algorithm implemented in the R library '*Rbeast'* (version 1.0.0, Zhao et al., 2019) to detect the places where the trend of inter-note interval changed within a call. This method allowed is to quantitatively define the point where calls transition from the introductory phase to the fast portion (i.e., when the slopes changes to zero). Then we counted the number of notes in the fast portion of the calls and divided this number by the duration of this section to compute the note repetition rate shown on the y-axis of panel (a). The x-axis of (a) correspond to the duration of the complete call, including the introductory section. Following Podos (1997), we split the call duration axis in bins of 4 seconds each (red vertical lines) starting from the individual with the shortest call duration. For each bin we identified the call with the highest note repetition rate (pink dots), and used these data to compute the upper-bound regression using ordinary least-squares regression (pink line and shading) (slope = -0.02, P = 0.0396, $r^2 = 0.43$). This upper-bound regression indicates that frogs cannot simultaneously maximize call duration and note repetition rate above the threshold value defined by the regression, and individuals close to the line have higher performance than those further away.

(b) Example of the Bayesian change point detection algorithm applied to the call shown in **Fig. 1b** of the main text. The top panel shows the raw inter-note interval data and the estimated trend as a black line. The middle panel shows the estimated slope of the trend, and the bottom panel shows the probability of the slope being negative (blue), positive (red) or zero (green). In this case, two break points were identified (vertical dashed lines, top panel), but only the second indicated the transition from the introductory section of the call to the fast note-repetition portion, where the trend slope is zero.

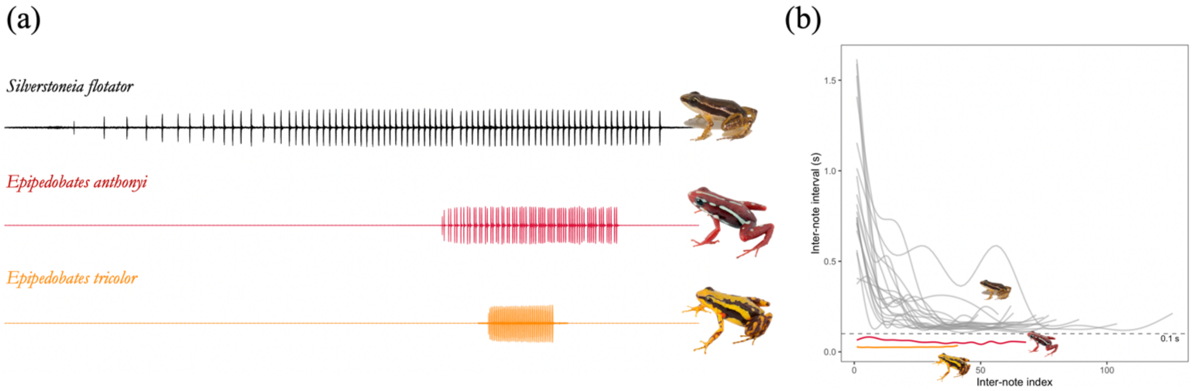


**Supplementary figure 3:** (a) Oscillograms of the advertisement vocalizations of *Silverstoneia flotator*, *Epipedobates anthonyi* and *Epipedobates tricolor*. All three calls are plotted on the same temporal scale. (b) Inter-note interval versus inter-note index of *S. flotator* (grey) compared to a call of *E. anthonyi* (red) and *E. tricolor* (yellow). The pictures of the frogs and vocalizations of *Epipedobates* were downloaded from the Anfibios del Ecuador website (https://bioweb.bio/faunaweb/amphibiaweb/).

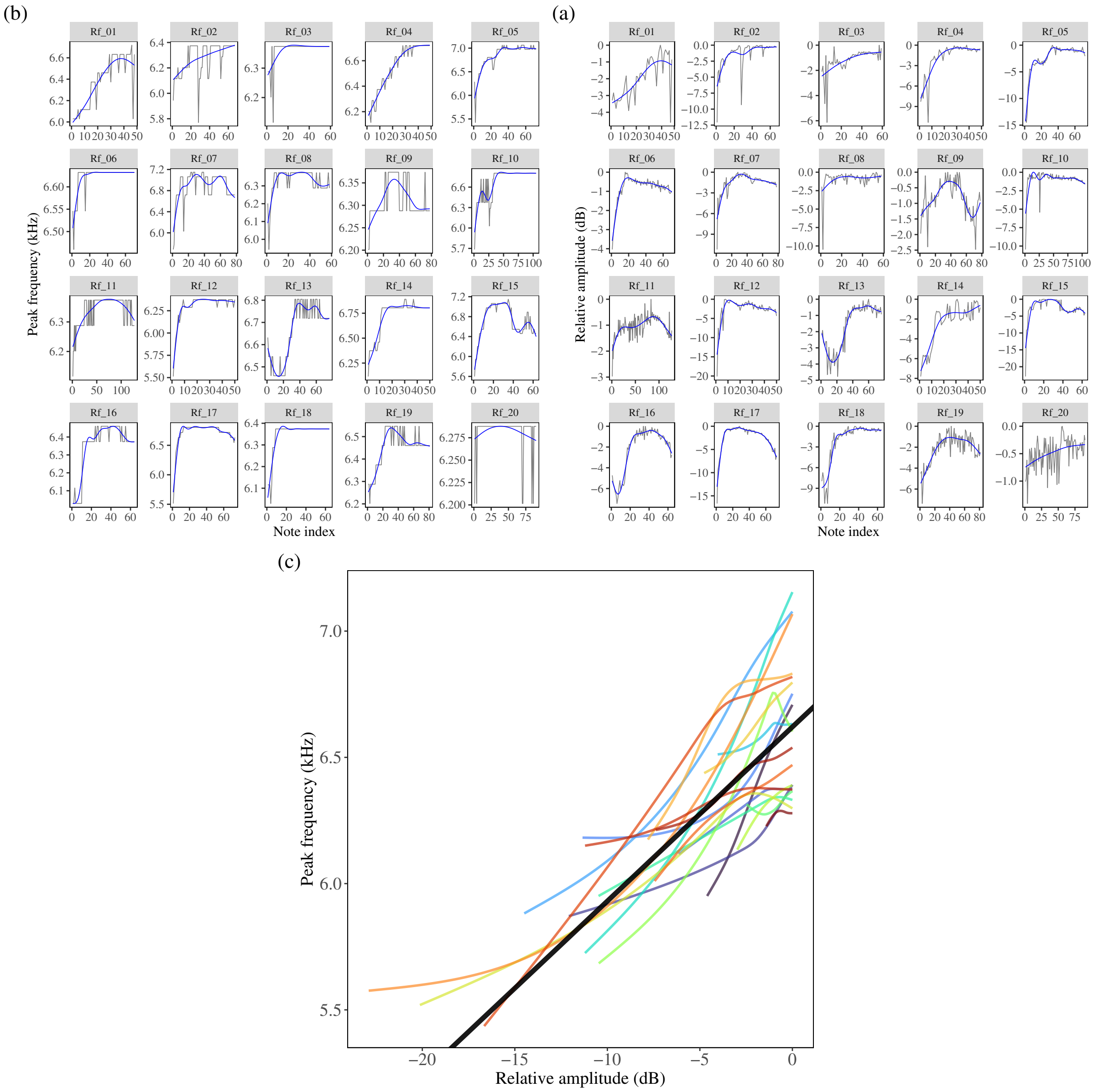


**Supplementary figure 4:** (a) Variation in peak frequency of the notes from the calls of twenty different rocket frog males. (b) Variation in the amplitude of the notes from the calls of the same twenty males as shown in panel (a). Note how note frequency and amplitude correlate across males, and how amplitude tend to be smaller at the beginning of the calls. Note index corresponds to the sequential progression of notes in a call, from the first note (note index = 1) to the last one. (c) Association between peak frequency and amplitude for the 20 males in panel (a) and (b). The coloured lines show the smoothed generalized additive function applied to each male, while the black line shows the estimate obtained from a linear-mixed effects models including each individual as a random component in the model. Amplitude was measured as the root-mean-square (RMS) amplitude of each note, and transformed to relative dB values with reference on the note with the highest RMS of each call.

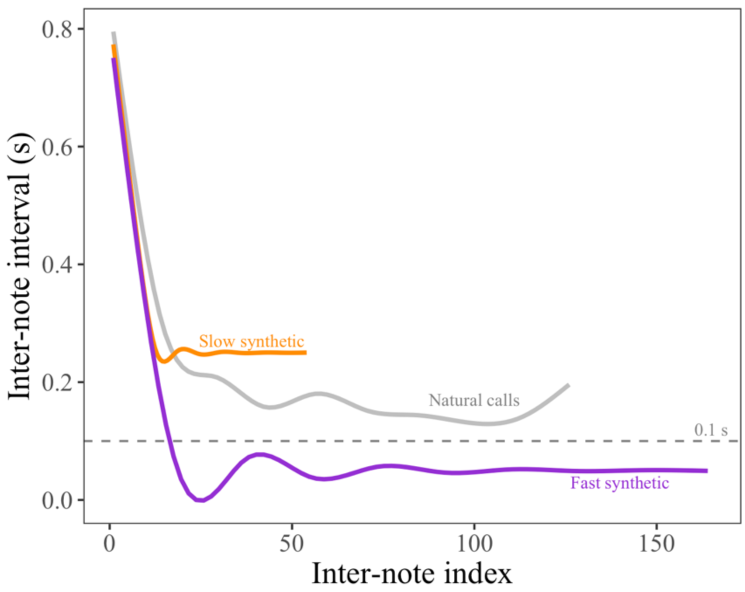


**Supplementary figure 5:** The inter-note intervals of the synthetic fast (orange) and slow (violet) stimuli as compared to the summary of the natural calls of 20 males (grey). The natural calls summarized are the same shown in **Fig. 1C** and **Supp. Fig. 1**. Inter-note intervals are smoothed with a generalized additive function for visual purposes.

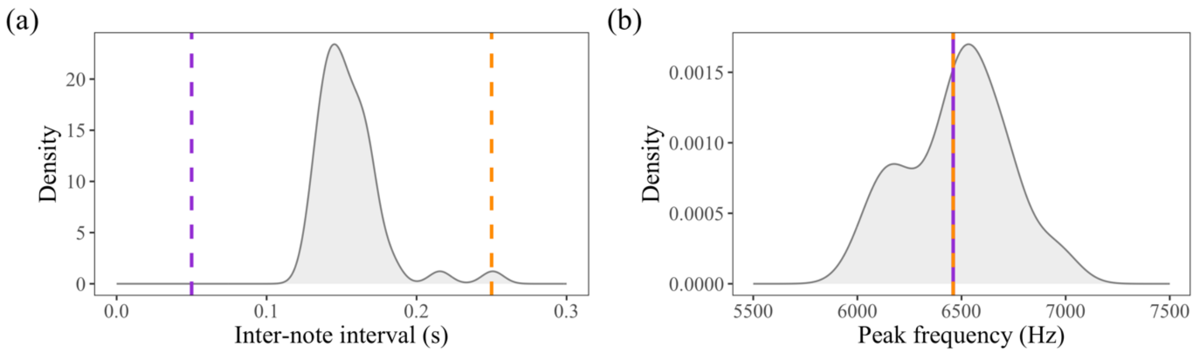


**Supplementary figure 6:** Comparison of (a) inter-note intervals and (b) peak frequencies of the N = 45 frogs tested in this study (grey kernel densities) relative to the synthetic stimuli. Fast (inter-note interval = 0.05 s, peak frequency 6460 Hz) and slow (inter-note interval = 0.25 s, peak frequency 6460 Hz) synthetic calls are depicted as purple and orange dashed lines, respectively.

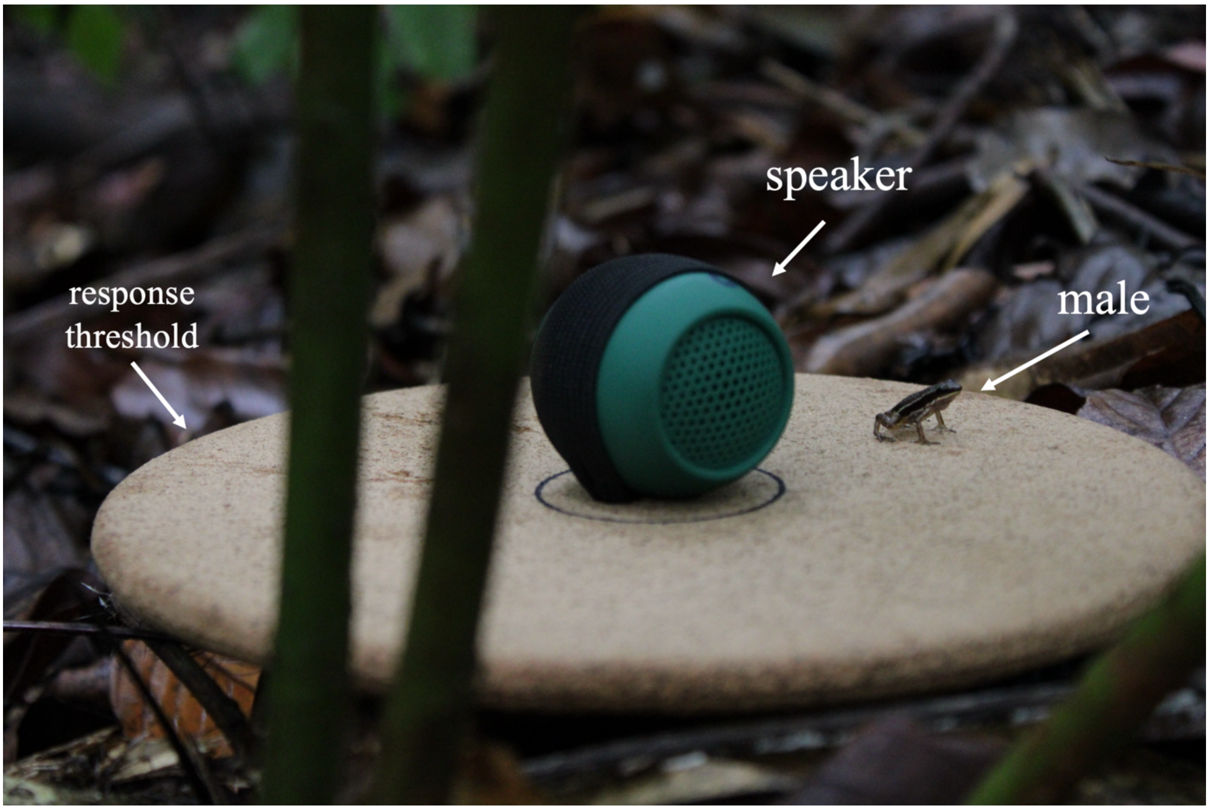


**Supplementary figure 7:** Example of an aggressive response given by a male. The response threshold is defined as males crossing the edge of the cork platform.

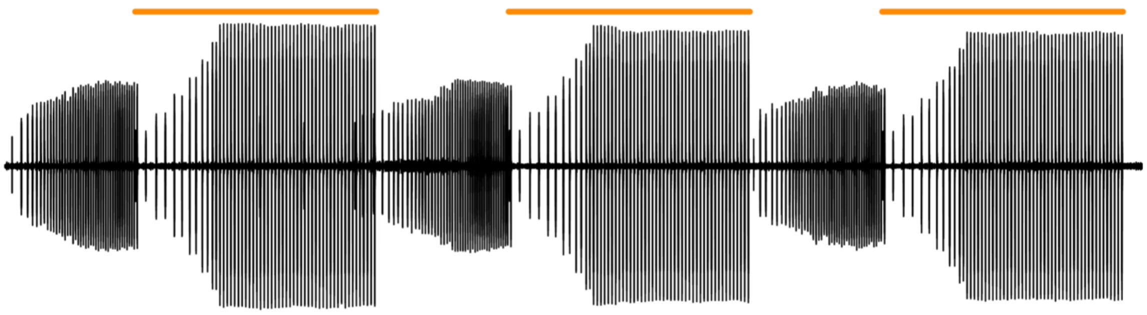

**Supplementary figure 8:** Example of the vocal responses of a male to three synthetic slow call. The duration of the synthetic call is show as a vertical orange bar on top of the oscillogram. This is one of the few examples of males that emitted complete calls during the 10-second silence between synthetic calls.

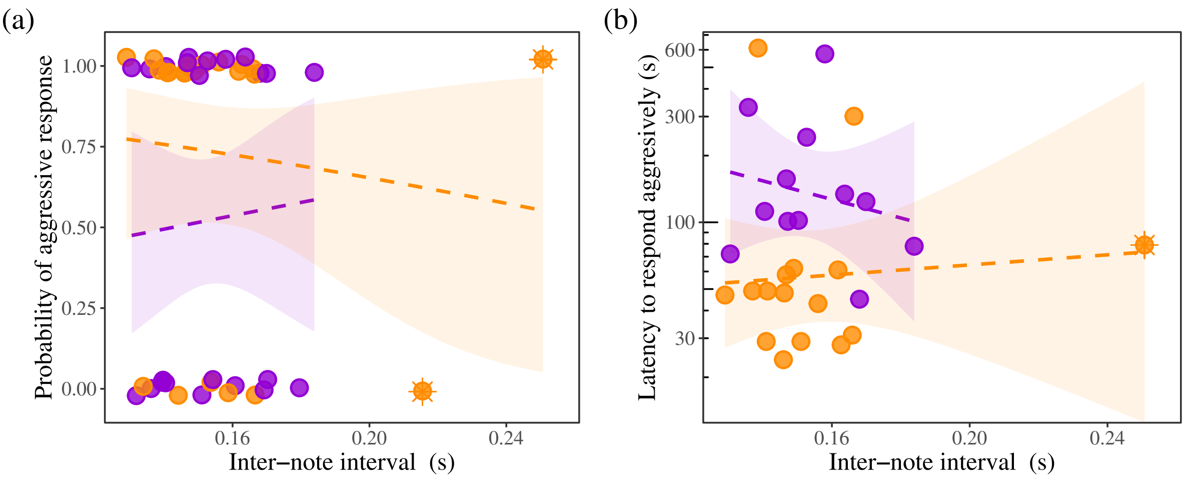


**Supplementary figure 9:** (a) Logistic and (b) linear regression analyses including the two individuals with unusually high (> 0.2 seconds) inter-note intervals marked as circle with an asterisk. The panel (a) corresponds to **Supp. Table 1**, and panel (b) to **Supp. Table 2**. Dashed lines indicate that slopes are not different from zero.

**Supplementary table 1:** Logistic regression without excluding outliers. See also **Supp. Fig. 6a**.

| Vocal trait | | Estimate | 95% CI | S.E. | z-value | P |
|---|---|---|---|---|---|---|
| Inter-note interval | Intercept | -1.187 | [-9.936, 7.278] | 4.276 | -0.278 | 0.781 |
| | Inter-note interval | 8.327 | [-46.701, 65.485] | 27.829 | 0.299 | 0.765 |
| | Stimulus:Slow | 3.494 | [-6.561, 13.708] | 5.060 | 0.690 | 0.490 |
| | Inter-note interval x Stimulus:Slow | -16.69 | [-82.311, 48.019] | 32.448 | -0.514 | 0.607 |

**Supplementary table 2:** Linear regression without excluding outliers. See also **Supp. Fig. 6b**.

| Vocal trait | | Estimate | 95% CI | S.E. | t-value | P |
|---|---|---|---|---|---|---|
| Inter-note interval | Intercept | 2.772 | [0.547, 4.997] | 1.078 | 2.572 | **0.017** |
| | Inter-note interval | -4.180 | [-18.567, 10.207] | 6.971 | -0.600 | 0.554 |
| | Stimulus:Slow | -1.192 | [-3.669, 1.285] | 1.200 | -0.993 | 0.330 |
| | Inter-note interval x Stimulus:Slow | 5.320 | [-10.636, 21.276] | 7.731 | 0.688 | 0.498 |

**Supplementary table 3:** Results from the linear mixed-model fitted to evaluate the vocal plasticity of males in response to the playback treatment.

| Fixed-effect | Sum. Squares | Mean squares | Num. df | Den. df | F-value | *P* |
|---|---|---|---|---|---|---|
| Period | 110373 | 110373 | 1 | 37 | 10.625 | **0.002** |
| Relative peak freq. | 342920 | 342920 | 1 | 37 | 33.010 | **< 0.001** |
| Stimulus | 19895 | 19895 | 1 | 37 | 1.915 | 0.175 |
| Period x Relative peak freq. | 143607 | 143607 | 1 | 37 | 13.824 | **0.001** |
| Period x Stimulus | 1333 | 1333 | 1 | 37 | 0.128 | 0.722 |
| Relative peak freq. x Stimulus | 25964 | 25964 | 1 | 37 | 2.499 | 0.122 |
| Period x Stimulus x Relative peak freq. | 1473 | 1473 | 1 | 37 | 0.142 | 0.709 |

Period: basal or playback.
Relative peak freq: higher or lower than the stimulus.
Stimulus: fast or slow synthetic call.

**Supplementary table 4:** Results of logistic regressions fitted to evaluate whether the probability of responding aggressively depends on the relative peak frequency of tested males (as either a continuous or categorical variable) and the stimulus.

| Vocal trait | | Estimate | 95% CI | S.E. | *z*-value | *P* |
|---|---|---|---|---|---|---|
| Relative peak frequency (continuous) | Intercept | 0.126 | [-0.803, 1.083] | 0.469 | -0.268 | 0.789 |
| | Relative peak frequency | -0.004 | [-0.009, 0.000] | 0.002 | -1.988 | 0.219 |
| | Stimulus:Slow | 0.857 | [-0.492, 2.289] | 0.697 | 1.229 | **0.047** |
| | Relative peak frequency x Stimulus:Slow | 0.008 | [0.002, 0.015] | 0.003 | 2.450 | **0.014** |
| Relative peak frequency (categorical) | Intercept | -0.588 | [-1.768, 0.475] | 0.558 | -1.054 | 0.292 |
| | Relative peak frequency:Lower | 1.841 | [0.048, 4.005] | 0.977 | 1.884 | 0.060 |
| | Stimulus:Slow | 2.293 | [0.573, 4.422] | 0.950 | 2.414 | **0.016** |
| | Relative peak frequency:Lower x Stimulus:Slow | -3.322 | [-6.298, -0.664] | 1.412 | -2.352 | **0.019** |

### Supplementary references

Zhao, K., Wulder, M. A., Hu, T., Bright, R., Wu, Q., Qin, H., ... & Brown, M. (2019). Detecting change-point, trend, and seasonality in satellite time series data to track abrupt changes and nonlinear dynamics: A Bayesian ensemble algorithm. *Remote Sensing of Environment*, *232*, 111181.
